# scCompass: An integrated cross-species scRNA-seq database for AI-ready

**DOI:** 10.1101/2024.11.12.623138

**Authors:** Pengfei Wang, Wenhao Liu, Jiajia Wang, Yana Liu, Pengjiang Li, Ping Xu, Wentao Cui, Ran Zhang, Qingqing Long, Zhilong Hu, Chen Fang, Jingxi Dong, Chunyang Zhang, Yan Chen, Chengrui Wang, Guole Liu, Hanyu Xie, Yiyang Zhang, Meng Xiao, Shubai Chen, The X-Compass Consortium, Yiqiang Chen, Ge Yang, Shihua Zhang, Zhen Meng, Xuezhi Wang, Guihai Feng, Xin Li, Yuanchun Zhou

## Abstract

Emerging single-cell sequencing technology has generated large amounts of data, allowing analysis of cellular dynamics and gene regulation at the single-cell resolution. Advances in artificial intelligence enhance life sciences research by delivering critical insights and optimizing data analysis processes. However, inconsistent data processing quality and standards remain to be a major challenge. Here we propose scCompass, which provides a data quality solution to build a large-scale, cross-species and model-friendly single-cell data collection. By applying standardized data pre-processing, scCompass integrates and curates transcriptomic data from 13 species and nearly 105 million single cells. Using this extensive dataset, we are able to archieve stable expression genes (SEGs) and organ-specific expression genes (OSGs) in human and mouse. We provide different scalable datasets that can be easily adapted for AI model training and the pretrained checkpoints with state-of-the-art (SOTA) single-cell foundataion models. In summary, the AI-readiness of scCompass, which combined with user-friendly data sharing, visualization and online analysis, greatly simplifies data access and exploitation for researchers in single cell biology(http://www.bdbe.cn/kun).

## Introduction

Cells are the fundamental units of life and a central focus of life sciences research^1, 2, 3^. High-throughput RNA sequencing enable to capture RNA in a large scale to reveal the whole genome-expression profiling^4, 5^. The emergence of single-cell RNA sequencing (scRNA-seq) can reveal the genetic structure and gene expression states of individual cells, reflecting cellular heterogeneity. The application of scRNA-seq has facilitated the generation and accumulation of vast amount of single cell omics data. This enables further insight into exploration of life processes and to understand cellular heterogeneity^6, 7^.

Extensive single cell datasets are essential for uncovering fundamental biological processes, transcriptomic foundation models offer potential to reveal complex interactions and regulatory mechanisms within organisms. The training of foundation models requires a large amount of high-quality transcriptomic datasets, like geneCompass^8^ (100 million cells), scGPT^9^ (80 million cells) and geneFormer^10^ (30 million cells). Their experimental results demonstrate that as data volumes increase, model performance correspondingly improves, emphasizing the urgent need for large, standardized, and model-friendly datasets. But the data integration and standardization pose great challenges. Platforms like CELLxGENE^11^, Human Cell Landscape (HCL)^12^, and Tabula Muris^13^ has improved the usability and integration of multiple datasets. CELLxGENE explores data to understand the mechanisms of human health. HCL is specialized in human gene expression, whereas Tabula Muris focues on mouse single cells. While these developments are promising, significant challenges still remains in terms of data consistency and accessibility. In particular, existing databases suffer from limitations such as limited organism, a lack of standardized procedures, inefficiencies and inaccuracies at downstream annotation processes, or inadequate quality control.

Therefore, we provide a single-cell dataset, scCompass, which provides a consistent process spanning human, mouse and other 11 species. We first gathered over 2 petabytes (PB) of raw single-cell RNA sequencing (scRNA-seq) data from public resources. Next by employing a unified data analysis and quality control (QC) process, we constructed detailed single-cell atlases for each species. These datasets consist of nearly 105 million single-cell transcriptomes. Leveraging this large-scale single-cell dataset, we aim to analyze and extract biological insights that are difficult to uncover in smaller datasets^14, 15^. Traditional housekeeping genes (HKGs) have been identified through large-scale bulk gene expression analyses^16, 17^. In this study, we leveraged single-cell data to explore and identify a novel set of stable expression genes (SEGs) at the single-cell level. These genes offer a promising foundation for future reference gene evaluation in transcriptomic research. scCompass includes over 30 organs (organ: exclude immune cells and nervous system) from human and mouse. We identify genes with organ-specific expression gene (OSGs) according to metadata, offering a valuable resource for elucidating the functional characteristics of different organs.

As the need for AI-ready single-cell datasets is growing, providing high-dimensional data with rich cellular detail that has the potential to foundation model pretraining^18^. To maximize this potential, we have constructed a series of datasets across varying scales, each carefully prepared through median normalization, expression value ranking, and standardized processing of filtered single-cell expression matrices. These datasets are optimized for direct integration with SOTA single-cell foundation models, including scGPT^9^, GeneFormer^10^, and GeneCompass^8^, and the pretrained foundation models are provided to enable researchers diverse single-cell downstream tasks. Furthermore, the standardized dataset can also be utilized for fine-tuning. By developing high-quality, AI-ready dataset, we aim to provide a robust foundation for optimizing dataset construction, reducing inter-dataset heterogeneity, accelerating AI model iteration, enhancing model generalization, and supporting the advancement of AI-driven research in life sciences.

We developed scCompass, a high-quality, uniformly processed dataset, from which we identified SEGs and OSGs. This resource provides AI-ready training datasets of various scales tailored to different foundational models, along with pretrained models. Additionally, scCompass features an interactive, user-friendly platform for data retrieval and sharing, enhancing accessibility for research and application.

## Results

### Construction of scCompass, a unified, large-scale, cross-species single cell transcriptomic data

Using a structured curation model (Fig. 1a), we integrated a cross-species (spanning human, mouse and other 11 species) single cell transriptome datasets from public repositories NCBI-GEO^19^, EMBL-EBI ArrayExpress^20^, and CNCB^21^. We first downloaded and analyzed all raw data that passed our screening, that is approximately 113 million expression profiles derived from a total of 15,337 samples. After QC, 104,637,783 cells remains, effectively filtering out less than 10% of cells in each species. Specifically, there are around 50 million cells each for human and mouse. (Fig. 1b, Fig. S1a). By observing the distribution of cell numbers across organs (Fig. 1c), we found that human and mouse have the most comprehensive coverage of organ types. This allows us to perform further analyses, including those of the OSGs.

**Fig. 1.**
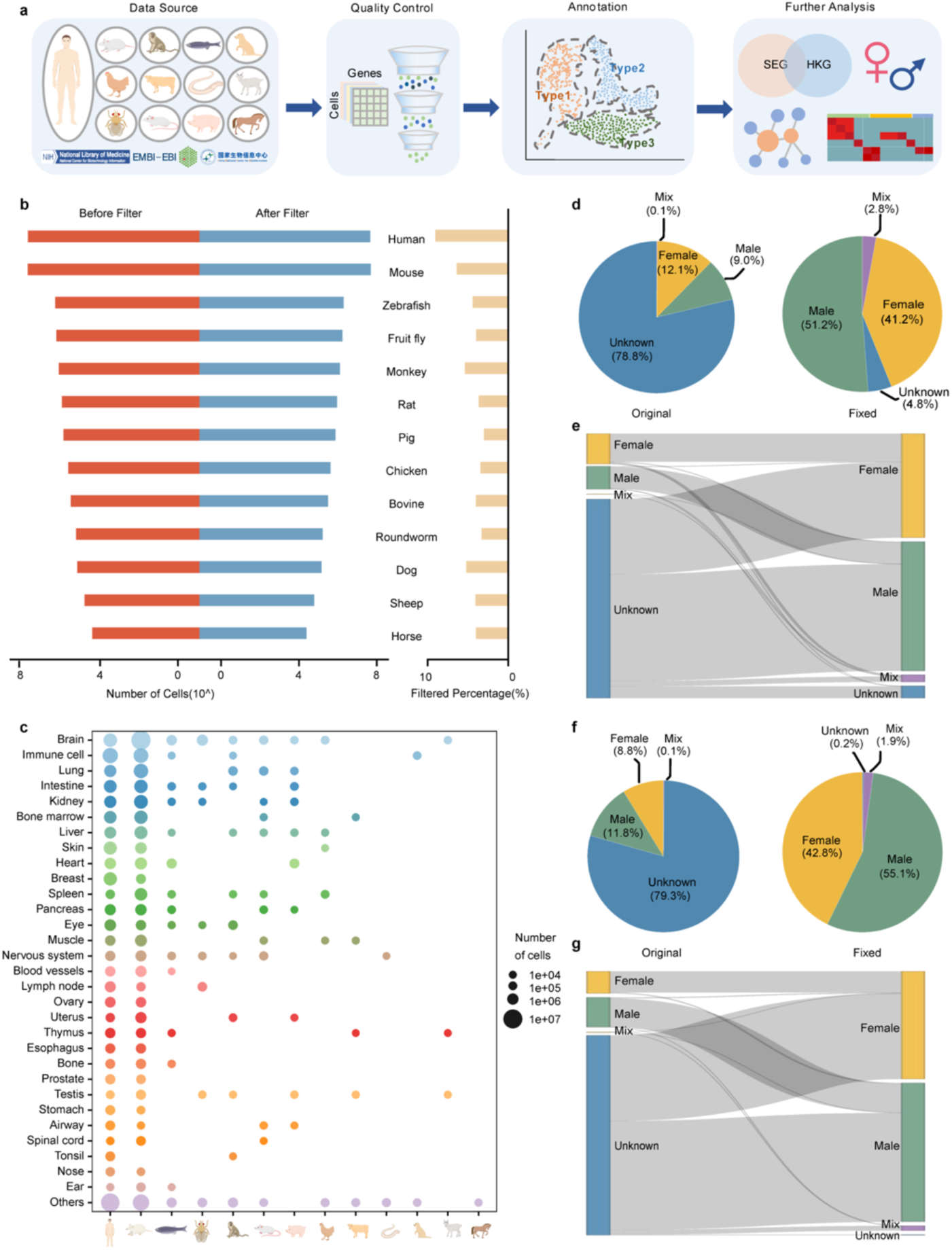
Data curation pipeline of scCompass. **a,** Illustration of scCompass data collection, QC, cell type annotation, and downstream analysis. **b,** Cell counts for 13 species before and after QC (left panel), along with the percentage of filtered out cells (right panel). **c,** Bubble chart depicting the cell proportions of 13 species across the top SI0 tissues. **d-g,** Sex correction in human and mouse. **d,** Human sex distribution before (left panel) and after correction (right panel). **e,** The Sankey diagram illustrates the results of sex correction for each human sample: left, original label, right, corrected label. **f,** Mouse sex distribution before (left panel) and after correction (right panel). **g,** The Sankey diagram illustrates the results of sex correction for each mouse sample: left, original label, right, corrected label.

We then meticulously annotated each dataset with species, sex, tissue, and cancer/normal distinction (Fig. 1d,f, and Fig. S1b). We observed that the sex information for a majority of human and mouse samples was labeled as “unknown”, with 78.8% of human samples and 79.3% of mouse samples lacking explicit sex designation. To address this, we defined X-Ratio and Y-Ratio thresholds (see Materials and Methods) by examing at the true distribution of X and Y chromosome gene expression in sex-specific tissues such as testis and ovary (Fig. S1c,d,e,f). Using these thresholds, we were able to assign sex attributes to the majority of samples previously labeled as “unknown” and verify the accuracy of existing sex labels. As a result, the proportion of samples with “unknown” sex attributes significantly decreased to 7.3% in the human dataset and 2.1% in the mouse dataset.

Overall, we construct a standardized and unified cross-species dataset, scCompass, following a structured curation process and provide a sex correction method.

### Constructing single-cell atlas of scCompass

Cell type annotation is essential to understand cellular heterogeneity. Our curated large-scale datasets display considerable variability in both annotation methods and quality. Therefore, we implemented a consistent cell type annotation process and create basic landscape of scCompass.

First, we performed a QC process on scCompass to only include high-quality cells. Initially, cells with fewer than 200 detected genes or samples with fewer than 4 cells were excluded. Subsequently, we removed cells expressing fewer than 7 protein-coding genes and mitochondrial gene expression accounted for over 15% in total. Due to the large volume of scCompass, it is impractical to label the cells with traditional manual annotation methods, we utilized cell type annotation tools SCimilarity^22^ and scMayoMap^23^, totally more than 200 cell types were annotated, covering the majority cell types (Fig. 2a).

**Fig. 2.**
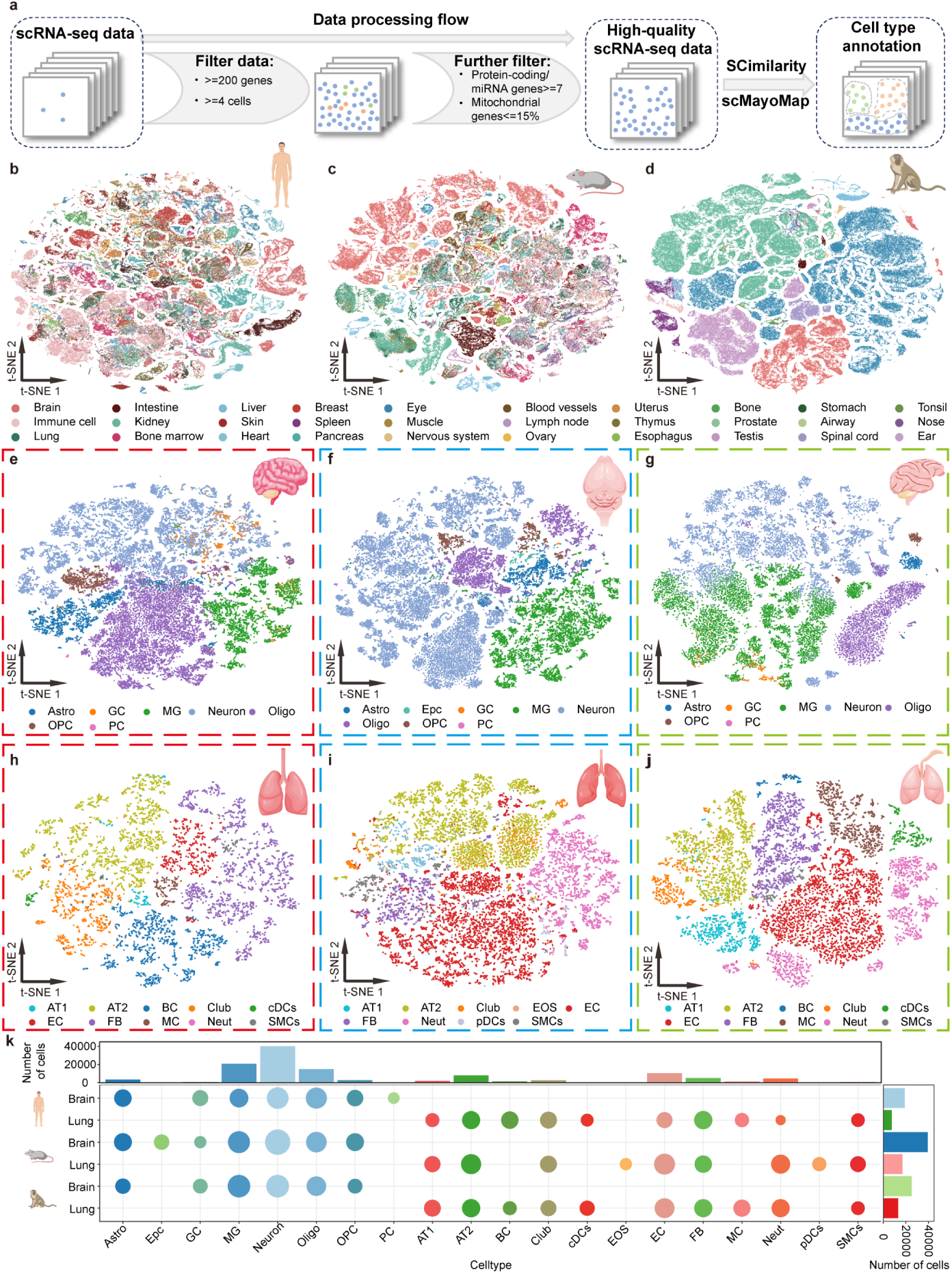
Single-cell atlas constrution of human, mouse, and monkey. **a,** Cell QC and cell type annotation (SCimilarity and scMayoMap). **b-d,** Single-cell atlas of human, mouse and monkey with t-SNE, colored dots represent different tissues. **e-g,** Cell types in brain of human (**e**), mouse (**f**), and monkey (**g**), colored dots represent different cell types. **h-j,** Cell types in lung of human (**h**), mouse (**i**), and monkey (**j**), colored dots represent different cell types. **k**, Statistics of the proportion and quantity of cell types in the brain and lung for human, mouse, and monkey. Astro, Astrocyte. Epc, Ependymal Cell. GC, Glial Cell. MG, Microglial Cell. Oligo, Oligodendrocyte. OPC, Oligodendrocyte Precursor Cell. PC, Pericyte. AT1, Type I Pneumocyte. AT2, Type II Pneumocyte. BC, Basal Cell. cDCs, Conventional Dendritic Cell. EOS, Eosinophil. EC, Endothelial Cell. FB, Fibroblast. MC, Mast Cell. Neut, Neutrophil. pDCs, Plasmacytoid Dendritic Cell. SMCs, Smooth Muscle Cell.

Then we illustrate the atlas of randomly select single cells (297,403 human cells, 297,324 mouse cells, and 292,131 monkey cells) across the top 30 tissues (Fig. 2b,c,d, for other species Fig. S2). We focused on the brain and lung, the top two organs in terms of cell numbers. We present the cell types identified in the brain tissues of human, mouse, and monkey, with cell counts of 37,423, 39,368, and 25,166, respectively. The main cell types observed include oligodendrocytes, oligodendrocyte precursor cells, microglial cells, astrocytes, glutamatergic neurons, neural cells, etc. (Fig. 2e,f,g). Similarly, we display the lung tissue cell distributions for human, mouse, and monkey, comprising 7,385, 16,887, and 13,025 cells, respectively. The major cell types in lung tissues were identified as alveolar type I (AT1) cells, alveolar type II (AT2) cells, and endothelial cells, etc. (Fig. 2h,i,j). The annotation results indicate that the major cell types for each organ are well-represented.

Finally, we illustrate the hierarchical clustering of different cell types in brain and lung (Fig. 2k). In brain, neuron, MG, and oligo are abundant, displaying similar distribution among species, highlighting their crucial roles in neural function and maintenance. Likewise, in the lung, AT2 and EC were predominant, with consistent proportions across species. These findings highlight the presence of key functional cell types in the brain and lung and underscore the cross-species conservation of those cell types.

Overall, we construct the single-cell atlas for scCompass. Our cell type annotation covers all major cell types in species.

### Identification of SEGs from large datasets

Unlike traditional bulk RNA sequencing data, scRNA-seq provides insights into significant transcriptional differences at the individual cell level^24, 25^. On the cell population level, a subset of genes known as HKGs consistently display stable expression across various tissues^17, 26^. Therefore, identifing SEGs within a large scale single cell data is essential for basic cellular function analysis.

Gene expression patterns differ across tissues, we quantified the distribution of zeros per gene at the single-cell level in human and mouse. Most genes exhibit zero expression in 80% of cells, indicating that ranking by zero-value rates effectively reflects gene stability across different cells (Fig. 3a,c). Then, we selected the same number of ranked genes according to the human and mouse HKG lists, named SEGs (Fig. 3b,d). We found that SEGs and HKGs share 52.6% overlap in human and 49.5% in mouse, suggesting intersection but also distinctiveness of SEGs.

**Fig. 3.**
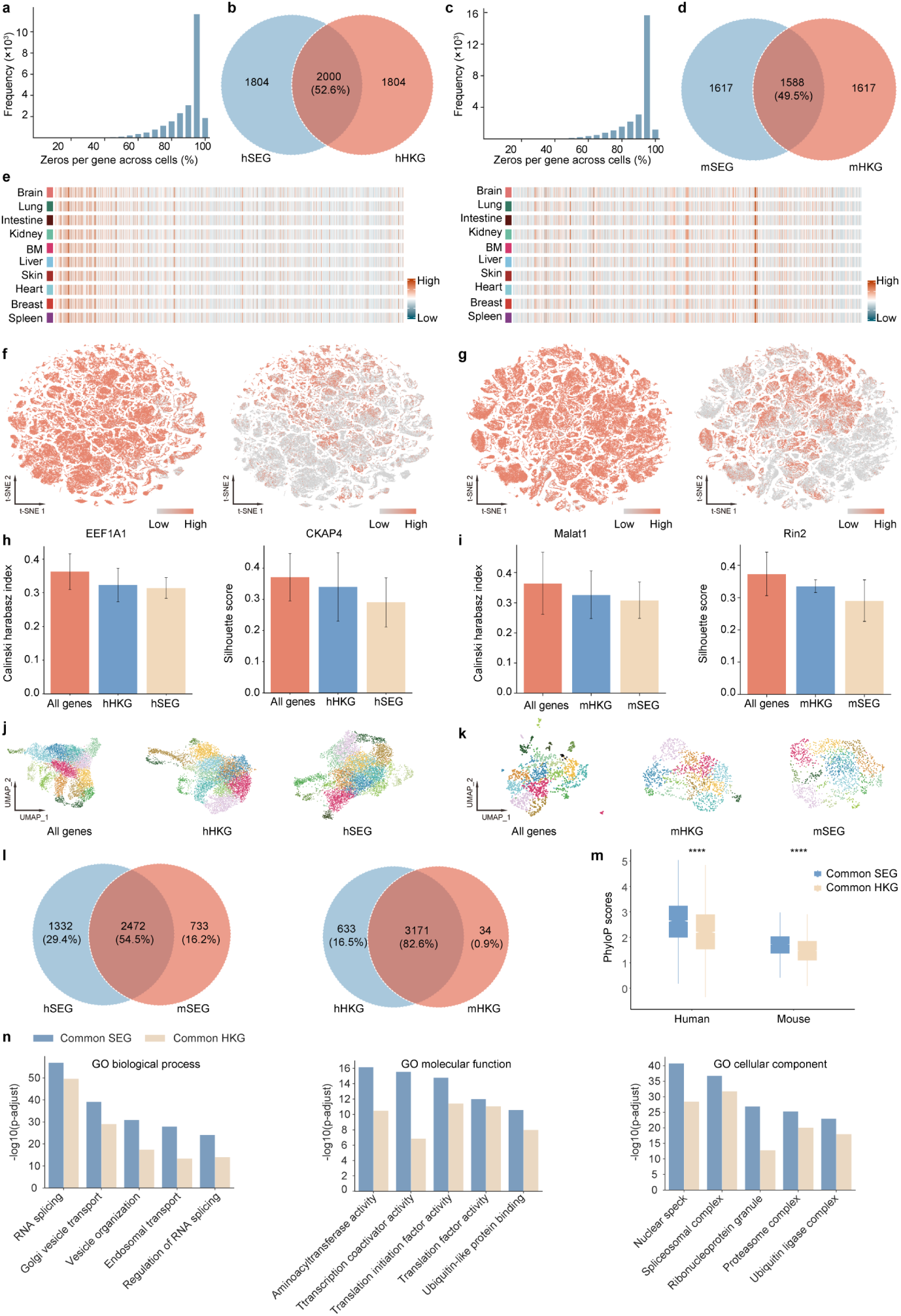
SEG analysis of human and mouse. **a,** The zeros per gene across human single-cell data. **b,** Number of common genes between SEGs and HKGs for human with venn plot. **c,** The zeros per gene across mouse data. **d,** Number of shared genes between SEGs and HKGs for mouse. **e,** Heatmap showing the distribution of SEGs in the top 10 organs for human (left panel) and mouse (right panel). **f.** tSNE visualization showing the expression pattern of EEF1A1 of unique hSEGs and CKAP4 of unique hSEGs, respectively, in indicated the top30 organs of human. **g,** tSNE visualization showing the expression pattern of Malat1 of unique mSEGs and Rin2 of unique mSEGs, respectively, in indicated the top30 organs of mouse. **h,i,** K-means clustering of 20 randomly sampled single-cell data evaluated with the Calinski-Harabasz index and silhouette score, using all expressed genes, HKGs and SEGs identified from this study for human (hSEGs) and mouse (mSEGs). **j,k,** UMAP plots generated from human (j) and mouse (k) single-cell data randomly selected from 20 samples using all expressed genes, HKGs, or SEGs. **l,** Common SEGs and HKGs between human and mouse. **m,** Comparison of conservation for common SEGs and HKGs in human and mouse genomes, with p-values calculated using a two-sided Wilcoxon rank-sum test. **n,** Overrepresentation analysis of SEGs shared between hSEG and mSEG (common SEGs) and HKGs shared between hHKG and mHKG (common HKG), using Gene Ontology (GO). (BM, Bone Marrow)

To demonstrate stable expression across different organs, we investigated SEG and HKG expression levels in pseudo-bulk data across top 10 organs for both human and mouse, they both displaying stable expression (Fig. 3e, Fig. S3a,b). The unique SEGs expression pattern are more stable than HKGs at single-cell resolution. For example, unique SEGs (EEF1A1, Malat1) showed pan-expression pattern while HKGs (CKAP4, Rin2) showed imbalanced expression in top 30 tissues (Fig. 3f,g and Fig. S3c,d).

To further validate the stability of SEGs, we evaluate the performance of all gene, HKGs and SEGs at clustering using Calinski harabasz index^27^ and Sihouette score^28^. The results showed that clustering with all genes performed best, followed by HKGs, with SEGs yielding the lowest performance (Fig. 3h,i). The UMAP visualizations (Fig. 3j,k) illustrate that cell clusters become less distinct from all genes to SEGs. This finding suggests that while SEGs have higher expression stability, they contribute less to the variance needed for effective clustering, as clustering algorithms rely on genes with variable expression to distinguish cell types.

Finally, we analyzed the evolutionary conservation of common SEGs (genes overlap between hSEGs and mSEGs) and common HKGs (genes overlap between hSEGs and mSEGs). The phyloP scores reveal that common SEGs exhibit significantly higher conservation (Fig. 3m). This suggests that SEGs may play fundamental roles during evolution. Subsequent Gene Ontology (GO) enrichment analyses (Fig. 3n) demonstrated that common SEGs are more enriched across GO terms compared to commonly defined HKG.

With the high-resolution scCompass dataset, we identified SEGs at the single-cell level that exhibit exceptional stability and pronounced evolutionary conservation.

### Defination of organ-specific expression genes

OSGs offer key insights into the specialized functions unique to each organ. The scCompass dataset covers 37 diverse organs, enabling a comprehensive and systematic identification of OSGs across a broad spectrum of organs (Fig. 4a,b).

**Fig. 4.**
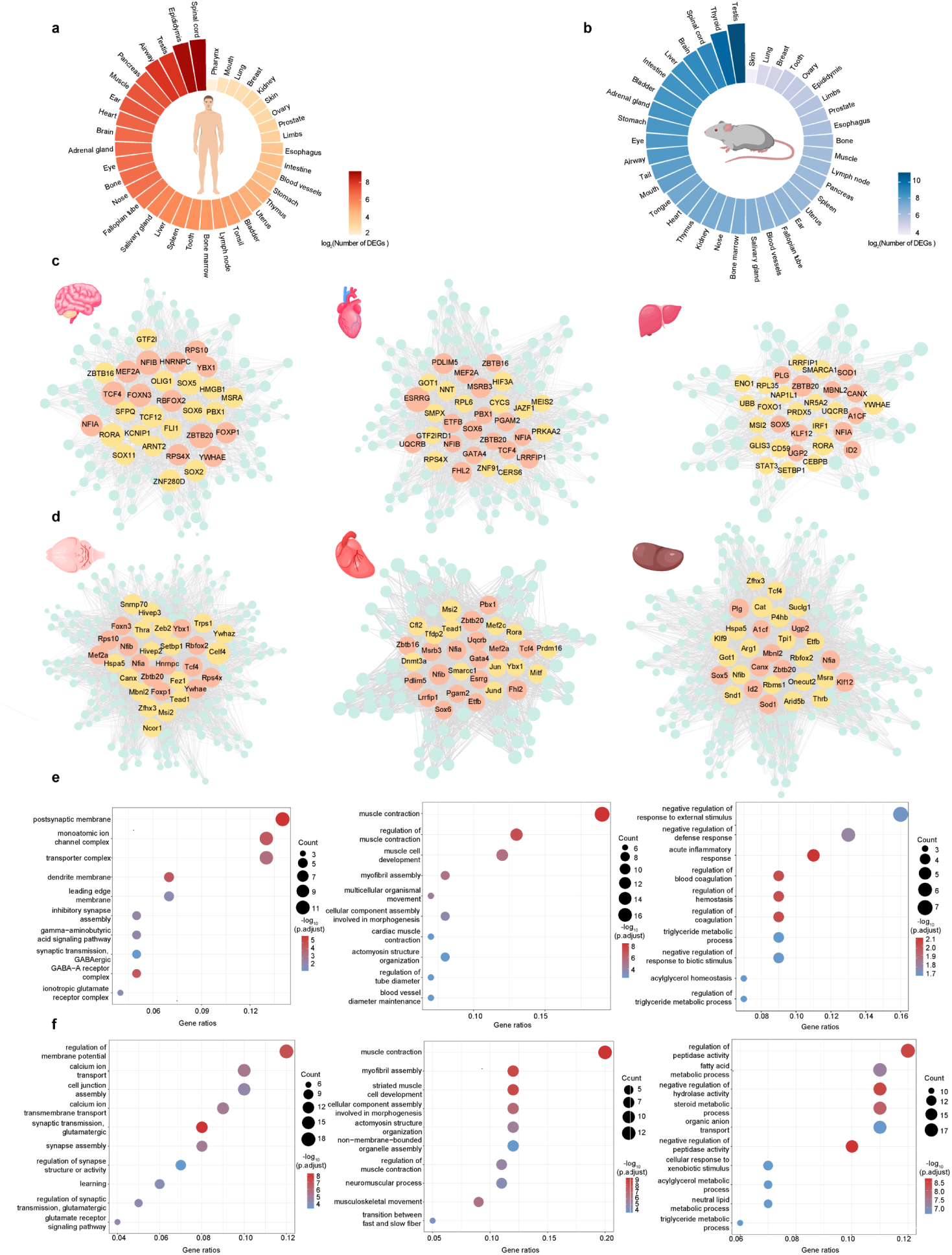
Organ differential expression gene analysis of human and mouse. **a,b,** Log values of differential expression genes in various organs between human and mouse. **c,d,** GRN constructed with OSGs of brain (left panel), heart (middle panle) and liver (right panel). Nodes with orange color represent the common TFs between human and mouse, the yellow nodes represent the distinct TFs of human or mouse, and the blue nodes represents the OSGs. **e,f,** GO enrichment related to cellular functions of predicted target genes in brain, heart and lung. Dot color represents significance level of enrichment analysis and dot size is count of target genes classified in GO terms. P-values were calculated using hypergeometric test. Multiple comparisons adjustment was performed using Benjamini and Hochberg method.

To explore the regulatory mechanisms underlying these OSGs, we constructed organ-specific GRNs of three vital functional organs (brain, heart, and liver) (Fig. 4c,d). Specifically, we identified 13 common TFs in the brain, 17 in the heart, and 11 in the liver. Several regulators for human brain (NFIB^29^ and OLIG1^30^), human heart (PDLIM5^31^ and FHL2^32^), human liver (FOXO1^33^ and ID2^34^), mouse brain (Msi2^35^ and Nfib^36^), mouse heart (Pdlim5^37^ and Tead1^38^), and mouse liver (Id2^39^ and Onecut2^40^), have been reported in the literature that is critically important in organs.

To elucidate the biological functions of these OSGs, we conducted GO analyses for the three organs (Fig. 4e,f). The enriched functions corresponded closely with the physiological roles of each organ. Specifically, in the human brain, OSGs were enriched in functions such as postsynaptic membrane and monoatomic ion channel complex, and in the mouse brain, they were associated with regulation of membrane potential and calcium ion transport. The muscle contraction is enriched both in the human and mouse heart. The human liver’s OSGs were involved in regulation of blood coagulation and triglyceride metabolic processes, while in the mouse liver, they were enriched in fatty acid metabolic processes and steroid metabolic processes.

Using the comprehensive organ metadata from scCompass, we analyzed the gene expression matrix to systematically identify OSGs for each organ.

### Evaluation of scCompass for AI-ready applications

Current single-cell transcriptomic foundation models require different input data formats: scGPT uses only single-cell expression matrices, GeneFormer relies on ranked gene expression sequences, and GeneCompass requires both expression matrices and ranked sequences. This inconsistency complicates standardized model evaluation and hinders broader applicability across diverse downstream tasks. To evaluate the adaptability of scCompass data in training single-cell AI models, we sampled 5 million human and 5 million mouse cells from both the scCompass and CELLxGENE. We quantified the gene numbers with detected expression values in each cell (Fig. 5a, b). For lower gene expression intervals, the cumulative frequency of CELLxGENE human data exceeded that of scCompass, suggesting that scCompass contains fewer missing values and higher data quality. This trend persisted in higher gene expression intervals, further affirming the superior data integrity of scCompass.

**Fig. 5.**
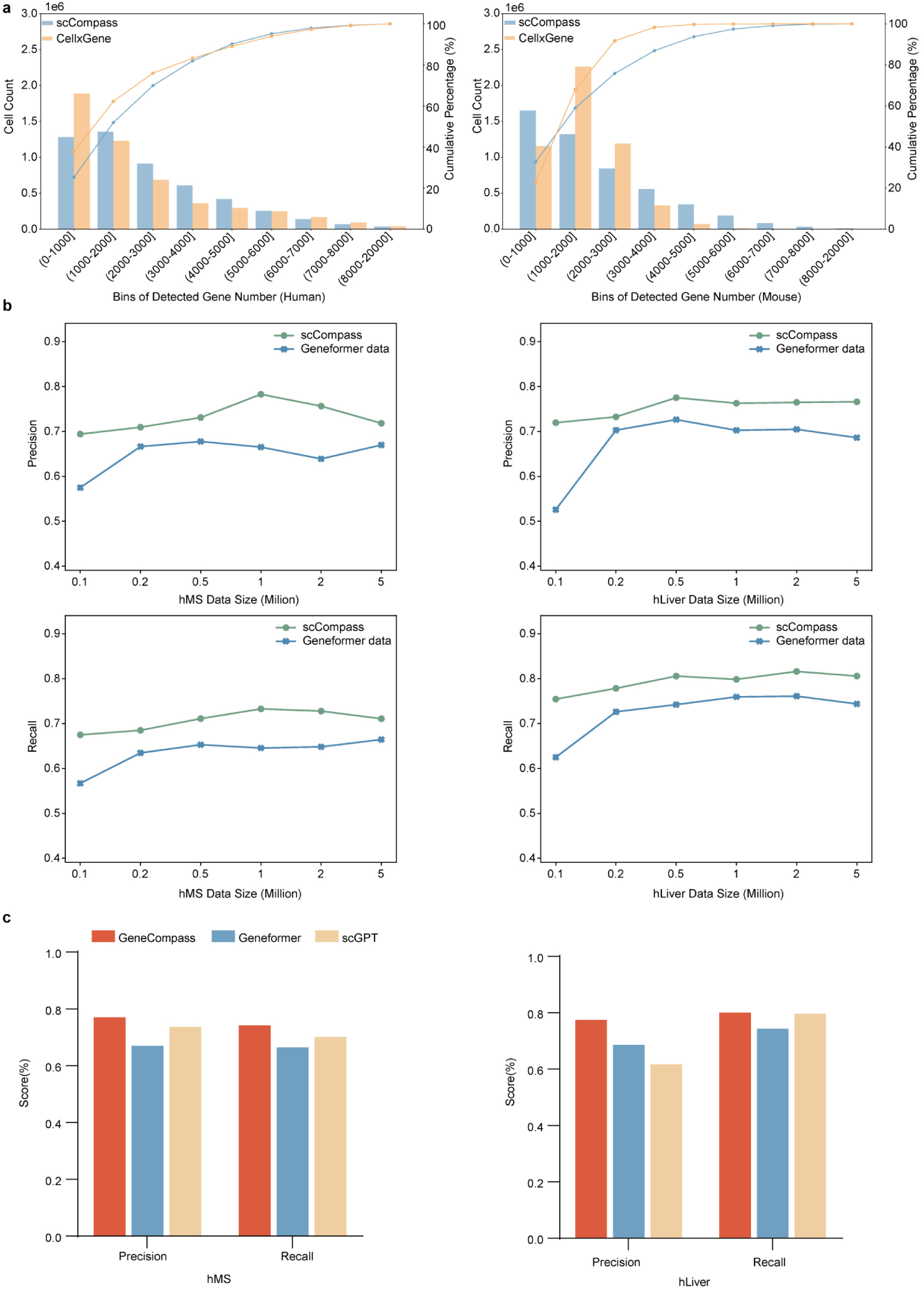
Evaluation of scCompass AI-Ready adaptability. **a,** A Pareto chart illustrates the gene expression profiles of cells sampled from scCompass and CELLxGENE. **b,** Precision and recall of GeneFormer models trained on various scales of single-cell samples, evaluated on hMS and hLiver for cell type annotation. Line plots, with green points representing scCompass pretraining and blue points for Geneformer. **c,** Precision and recall evaluation of cell type annotation tasks for different models (scGPT, GeneFormer, GeneCompass) on 5 million human sampled cells.

We pretrained scGPT on various scales of the scCompass and Geneformer datasets from CELLxGENE and evaluated model performance in cell annotation tasks (Fig. 5c, Fig. S4). scCompass consistently yielded higher recall and precision, with a more pronounced advantage at larger dataset scales. We also present the cell type annotation results for the 5 million cells in scCompass with pretraining on different foundation models (scGPT, Geneformer, and GeneCompass) (Fig. 5d). The results demonstrate precision and recall values consistently above 0.6, with peak values approaching 0.8, confirming the effectiveness of our scCompass dataset across the three AI models.

We constructed scalable, AI-ready datasets from scCompass and performed pretraining on SOTA single-cell foundation models. Using standardized cell type annotation tasks, we conducted a unified evaluation across all models.

### Interactive system for single-cell transcriptome data sharing and analysis

We provide a user-friendly interface incorporating search, browse, statistics, download, analysis and model features for scCompass exploration (http://www.bdbe.cn/kun). The system allows users to search and filter datasets, and provides interactive visualization of cells and gene expression pattern (Fig. 6a,b). Statistics for the entire dataset are provided, categorized by species and organ type (Fig. 6c). To enhance data accessibility, individual count matrix (h5ad format), metadata, and source code are downloadable (Fig. 6d). The series of online tools provides coding-free solutions for subsequent data annotation, normalization, and extension of the scCompass dataset (Fig. 6e). For AI-ready, the model feature include multi-scale, model-friendly datasets of varying sizes, as well as models and integrated embeddings (Fig. 6f).

**Fig. 6.**
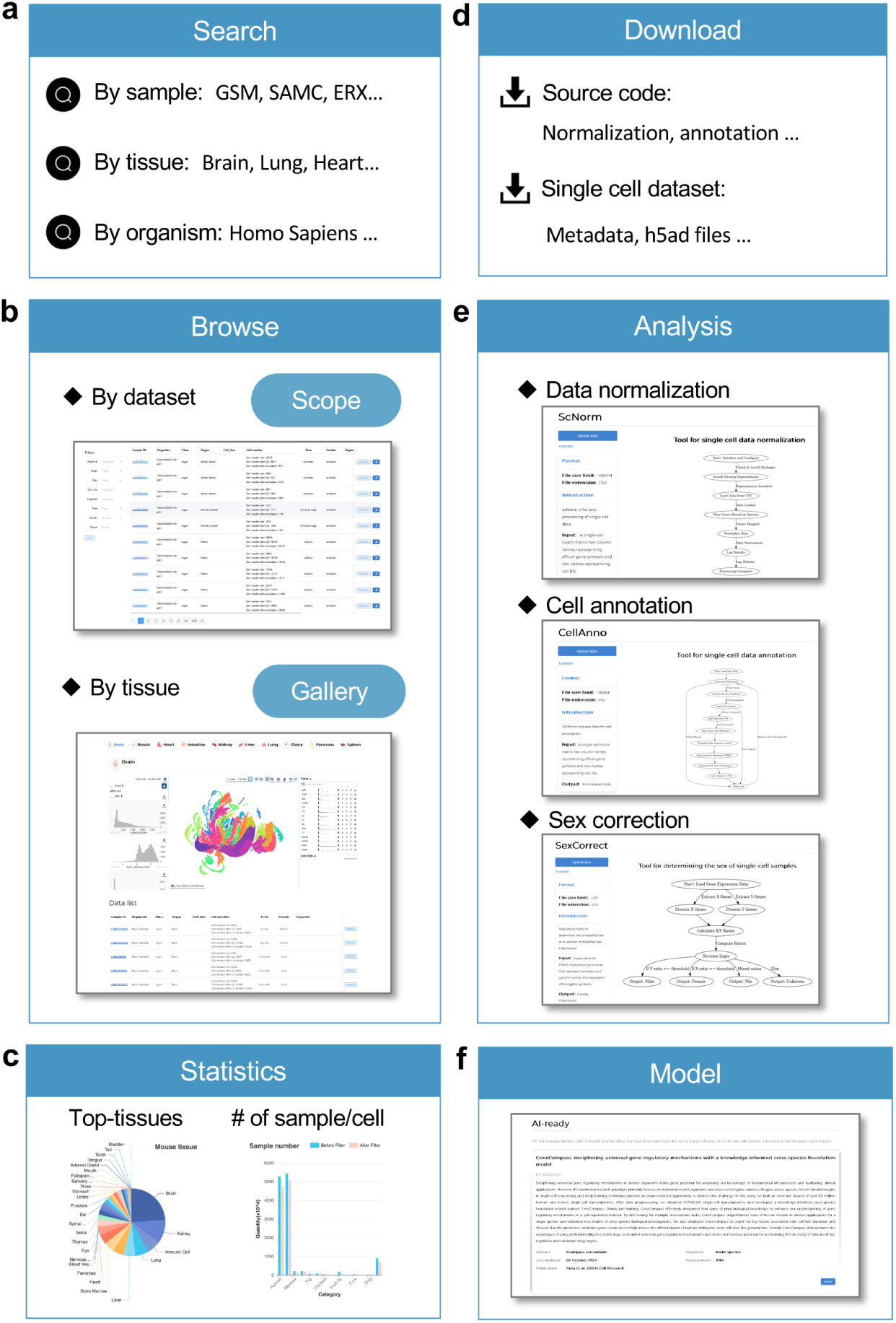
Illustration of different modules in the scCompass system. **a,** Search module accepts diverse search entries including sample ID, tissue, and organism. **b,** Interactive visualization in the Browse section for individual dataset of single-cell sample and integrated datasets of the same tissue. **c,** Statistics figures shows top tissues that have the most numbers of samples or cells across all datasets. **d,** Download feature provides source code and h5ad file for each sample. **e,** Analysis section includes three online tools scNorm, cellAnno and sexCorrect for data normalization, cell annotation and sex determination. **f,** The Model module contains models and embeddings that have been trained or fine-tuned with scCompass.

We developed an interactive single-cell transcriptome sharing and analysis system based on scCompass, offering data resource downloads and model-friendly evaluation datasets.

## Discussion

We proposed scCompass, an integrated cross-species scRNA-seq database for AI-ready. First we applied a rigorous, standardized QC process achieving 105 million high-quality single-cell transcriptomes, and we corrected the sex attribute of sample metadata. Then we utilized tools to annotate cell types, constructing single-cell atlases for each species. With this large-scale dataset, we identified SEGs and OSGs in both human and mouse, SEGs are highly consistant in different tissues, and can be used as reference genes in single cell resolution, and OSGs can offer valuable insights into organ functions. To facilitate the training of AI foundation models, we curated model-compatible datasets at multiple scales and provided comprehensive benchmarking results using several SOTA pretrained models. Lastly, we developed a multi-fuctional interactive data sharing and analysis system, enhancing accessibility and usability for research.

Currently, scCompass focuses exclusively on single-cell transcriptomic data, limiting its scope to a single modality. As multimodal omics^41^ analysis becomes increasingly important, there is a pressing need to expand beyond transcriptomics to incorporate additional data types, such as spatial transcriptomics and epigeomics. At present, we support rapid adaptation for single-cell foundation models. Future updates will include enhanced AI-ready features^42, 43, 44^, such as data-as-a-service (DaaS), model-as-a-service (MaaS), and a secure, unified evaluation framework using federated learning, aimed at enabling comprehensive, integrated, and scalable data analysis.

In conclusion, scCompass is a high-quality, shareable, AI-ready database through strict and consistent standardization processes. It enables researchers in the life sciences and artificial intelligence fields to access and utilize the largest single-cell database. The data sharing system provides new potential for large-scale biological insights and establishes essential infrastructure for the AI for Science research paradigm.

## Materials and Methods

### Multi-species data collection and preprocessing

We acquired 15,337 single-cell RNA sequencing (scRNA-seq) samples from 13 species, including human, mouse, monkey, roundworm, zebrafish, fruit fly, rat, pig, bovine, dog, horse, chicken, and sheep. These datasets were retrieved from various sources, including the Gene Expression Omnibus (GEO) of the National Center for Biotechnology Information (NCBI), European Molecular Biology Laboratory-European Bioinformatics Institute (EMBL-EBI) ArrayExpress, and the China National Center for Bioinformation (CNCB), using the keywords combinations such as “high throughput sequencing + 10X”, “RNA-seq + cellranger” and “10X RNA-seq”. After validating the integrity of the Sequence Read Archive (SRA) files, we used standardized tools, including sratoolkit^45^ (version 2.11.0) and pfastq-dump (version 0.1.6), for data processing. Once the SRA files were converted to fastq format, the resulting fastq files were analyzed with CellRanger^46^ (version 7.0.1) default parameters to generate gene expression matrices with the reference genome (human: hg38, mouse: mm10, monkey: Mul_10, rat: mRatBN7.2, bovine: ARS-UCD1.2, roundworm: WBcel235, dog: UU_Cfam_GSD_1.0, zebrafish: GRXzll, fruit fly: BDBP6.32, horse: EquCab3.0, chicken: gca000002315v5.GRCg6a, sheep: Oar_rambouillet_v1.0, pig: Sscrofall.1) for each species^47^, which were then used for downstream analysis.

### Quality control and data filtering

To ensure data quality and biological relevance, we applied stringent filtering at both the sample and cell levels. First, single-cell sequencing samples with fewer than 200 detected genes or fewer than 3 cells were excluded from further analysis. At the cellular level, we filtered out cells expressing more than 7 protein-coding or miRNA genes, and cells with a proportion of mitochondrial gene expression exceeding 15%. We also removed cells with an expressed gene count exceeding 3 standard deviations from the mean in each sample’s gene expression matrix, and excluded genes not found in the core gene list to prioritize high-quality data. After applying these QC measures, we retained 15,270 samples from 13 species, containing 104,637,783 single-cell datasets for further analysis.

### Cell type annotation

Due to the large scale and high dimensionality of single-cell data, manual cell type annotation is labor-intensive and subject to human bias. Thus, for human cell type annotation, we employed the Scimilarity^22^ tool, while for mouse data, we used ScMayoMap^23^ as the reference annotation database. For other 11 species, including zebrafish, we mapped homologous genes between each species and human, followed by cell type annotation using Scimilarity. Moreover, we dropped the low-quality annotated cells (cell numbers per annotated cell type less than 20).

### Sex correction analysis

We first calculated the proportion of chrX and chrY genes in sex-specific organs, namely the testis and ovary, from both human and mouse samples. The list of chrX and chrY genes is downloaded from Ensembl^47^. To determine sex, we defined the Y-Ratio and X-Ratio as follows:

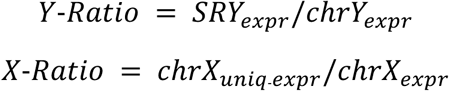

Here, SRYexpr refers to the expression of SRY gene (SRY^48^ for human, Sry^49^ for mouse), chrYexpr refers to the summary expression of chrY genes. chrXuniq-expr refers to the summary expression of unique genes of chrX, exclude homologous genes between chrX and chrY, chrXexpr refers to the summary expression of chrX genes.

We utilized the sex-specific organs, testis and ovary, to calculate Y-Ratio and X-Ratio values, establishing an initial threshold for sex determination. Using these initial thresholds, we performed sex predictions across different samples and compared them with existing sex labels. We then refined the thresholds based on incorrectly predicted samples, optimizing the final criteria for sex determination.

A human individual is determined as male if the Y-Ratio is ≥ 0.001188 or the X-Ratio ≤ 0.000070 with a Y-Ratio > 0.000106. Female classification requires an X-Ratio ≥ 0.000001 and Y-Ratio ≤ 0.000106. An X-Ratio > 0.000070 with a Y-Ratio between 0.001188 and 0.000106 indicates a “mix” chromosomal profile. Samples that did not meet these criteria were labeled as “unknown”.

A mouse individual is determined as male if the Y-Ratio is ≥ 0.000065 or the X-Ratio ≤ 0.000008 with a Y-Ratio between 0.000004 and 0.000065. Female classification requires an X-Ratio ≥ 0.000555 and Y-Ratio between 0.000004 and 0.000065, or X-Ratio > 0 with Y-Ratio ≤ 0.000004. An X-Ratio between 0.000008 and 0.000555 with a Y-Ratio between 0.000004 and 0.000065 indicates a “mix” chromosomal profile. Samples that did not meet these criteria were labeled as “unknown”.

### SEGs identification and assessment

We first quantified the proportion of zero counts for each gene across all cells in human and mouse samples by calculating the ratio of cells with zero counts to the total number of cells. Genes were ranked by their zero-rate values. Next, we performed pseudo-bulk processing on single-cell data and generate FPKM^50, 51^ (fragments per kilobase of transcript per million mapped reads) matrix from different species and calculated the CV^52, 53, 54^ for each gene as the ratio of the standard deviation to the mean expression. Genes were then ranked based on their CVs. We combined the rankings from the zero-rate and CV to generate a final ranking of gene stability.

To assess gene stability, we used HKGs as a reference^55^. We selected same number of HKGs from top-ranked stable genes, defining these as SEGs. To evaluate the expression stability of these gene sets across different cell types and biological systems, we pseudo-bulk processed single-cell data from ten organs (brain, lung, intestine, kidney, bone marrow, liver, skin, heart, breast, and spleen) and converted the data to FPKM. We visualized the expression patterns of SEGs and HKGs using heatmaps generated with the ComplexHeatmap^56^ R package.

Clustering was performed on each single-cell RNA sequencing dataset using the Seurat^57^ package, and clustering performance was evaluated using the Calinski-Harabasz index^27^ and Silhouette score^28^. We compared three scenarios: (i) using all expressed genes, (ii) using HKGs from bulk RNA-seq data, and (iii) using SEGs identified in this study. Clustering was repeated 20 times with random subsampling to account for variability. The SEGs and HKGs in human and mouse were replaced with their respective homologous genes, yielding a set of common SEGs and common HKGs. To characterize the evolutionary conservation of those common genes, we downloaded phyloP^58, 59^ scores for the human (hg38) and mouse (mm10) genomes from the UCSC Genome Browser^60^. Exonic bases were mapped based on GENCODE gene annotations, and a mean conservation score was calculated for each gene. We assessed the correlation between gene conservation using Pearson correlation coefficients^61^ and compared these features between common SEGs and common HKGs in both human and mouse.

We evaluated the overrepresentation of common SEGs and common HKGs by comparing each gene set to GO^62^ terms using the GO database with R package clusterProfiler^63^. Statistical significance was assessed using Fisher’s exact test, and the top enriched GO terms were merged for interpretation.

### Organ-specific OSGs definition

For organ-specific OSGs identification, we calculated the zero-rate of genes in single-cell data from different organs in both human and mouse datasets. We defined organ-specific genes as the propotion of zero values is no more than 90% of cells in a given organ but expressed in no more than two organs.

To enhance gene number and expression correlation in high-throughput single-cell mRNA datasets, we pooled data from 100 cells within the same cluster, and generate pseudo-cells to support genetic network analysis^64^.

Based on these pseudo-cell datasets, we used the tool pySCENIC^65^ to infer regulatory circuits and identified transcription factor-target gene (TF-TG) interactions for organ-specific genes. Only TF-TG interactions with the highest regulatory impact were retained for constructing regulatory networks (30 weights). These representative TF-TG networks were visualized using Cytoscape^66^ and the R package igraph^67^. Finally, we performed GO enrichment analysis on the organ-specific genes to uncover their functional roles within each organ.

### Evaluation of the Dataset Quality for AI-ready

To assess the quality of our dataset, we randomly selected 5 million cells each from geneCompass and CELLxGENE for both human and mouse. We initially quantified the gene numbers with non-zero expression values in each cell, then binned these counts in 1,000-gene intervals. To validate the quality of our constructed datasets, we randomly sampled single-cell data of varying scales from scCompass and Genecorpus-30M^10^ to build pre-training datasets and trained them on the specific foundational model, Geneformer. We then evaluated the model’s performance through downstream tasks involving cell type annotation. Initially, we created datasets of different data sizes 100 thousands, 200 thousands, 500 thousands, 1 million, 2 million, and 5 million. These datasets were used to pre-train three models: scGPT^9^, Geneformer^10^, and GeneCompass^8^. During the pre-training phase, we adhered to the hyperparameters specified in the original papers of these models, ensuring consistency in experimental setup.

To ensure a fair comparison, the pre-training process for each model involved keeping the hyperparameter settings consistent with those outlined in their respective publications, such as learning rate, batch size, and optimizer. We adjusted the number of epochs based on the dataset size, setting them at 10, 8, 7, 5, 4, and 3 for the datasets of 100 thousands, 200 thousands, 500 thousands, 1 million, 2 million, and 5 million cells, respectively. To further evaluate the performance of the trained models, we used the human multiple sclerosis (hMS)^8^, lung (hLung)^68^, and liver (hLiver)^69^ datasets as downstream tasks for cell annotation. The hyperparameter settings for each model in these downstream tasks were as follows: for GeneCompass, the learning rate was set to 5e-5, batch size to 16, and the number of training epochs to 50; for Geneformer, the learning rate was set to 5e-5, batch size to 16, and epochs to 50; and for scGPT, the learning rate was similarly set to 5e-5, batch size to 16, and epochs to 50.

Recall and precision were employed as evaluation metrics to assess the performance across the three datasets. Recall, also known as sensitivity, quantifies the proportion of true positive instances that are correctly identified by the model, providing a measure of completeness. The recall (R)^70^ is defined as:

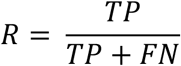

where TP represents True Positives, which are the number of correctly classified positive instances, and FN represents False Negatives, which are the number of positive instances incorrectly classified as negative.

Precision quantifies the proportion of correctly identified positive predictions among all instances predicted as positive, reflecting the accuracy of positive predictions. The precision (P)^70^ is given by:

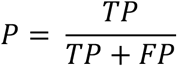

where FP represents False Positives, which are the negative instances incorrectly classified as positive by the model.

These metrics provided a comprehensive understanding of each model’s performance in terms of both identifying true positives effectively (Recall) and minimizing false positives (Precision). We evaluate the achieved models’ performance using two cell type annotation task datasets hMS and hLiver.

---

**Fig. S1.**
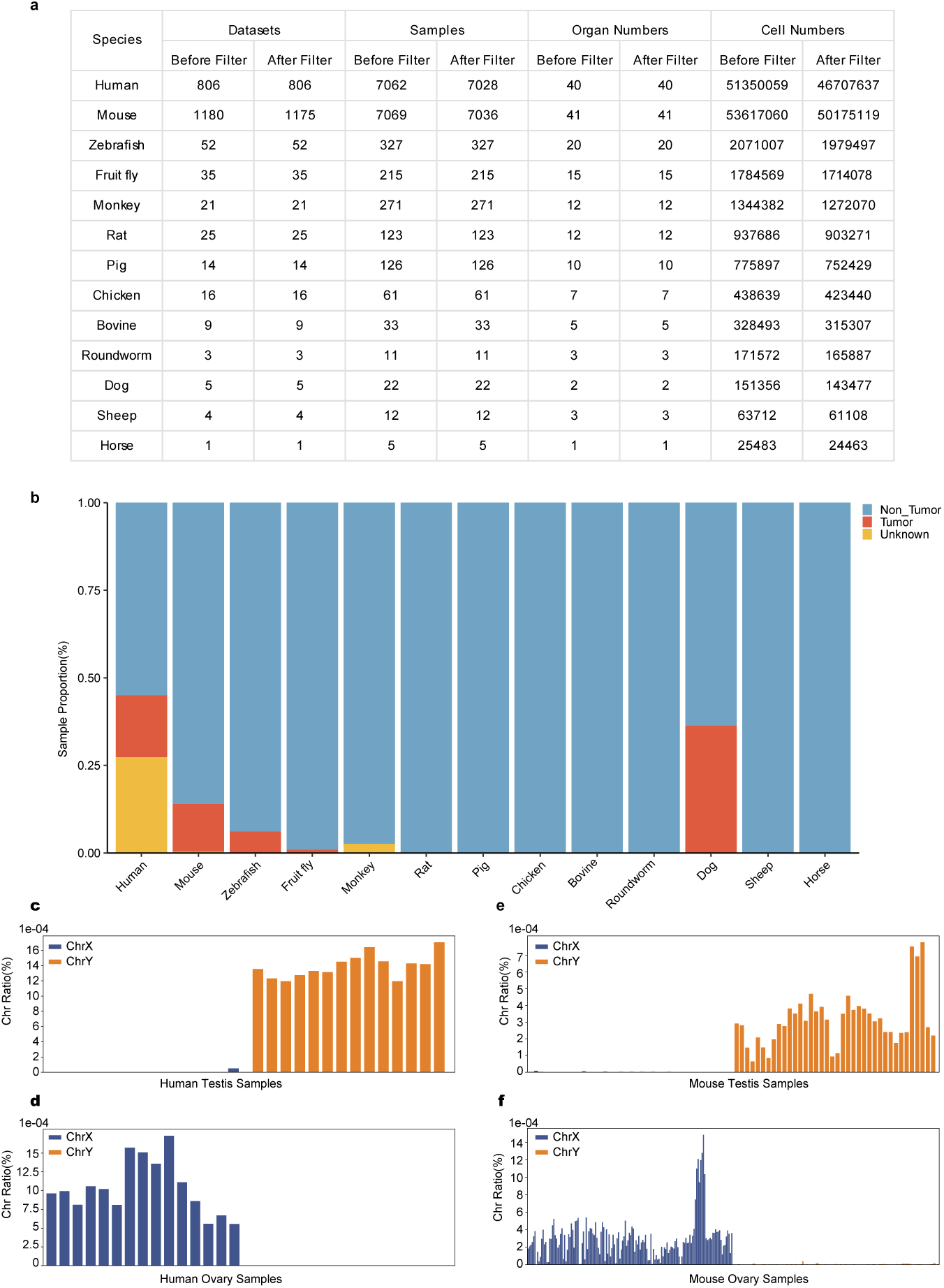
scCompass Statistical Analysis. **a,** Statistics of pre- and post-quality control results for datasets, samples, tissues, and cell counts across thirteen species in scCompass. **b,** Proportion of tumor samples across the thirteen species. **c,e,** Proportion distribution of X and Y chromosomes in human and mouse tesits, with X (blue) and Y (orange). **d,f,** Proportion distribution of X and Y chromosomes in human and mouse ovaries, with X (blue) and Y (orange).

**Fig. S2.**
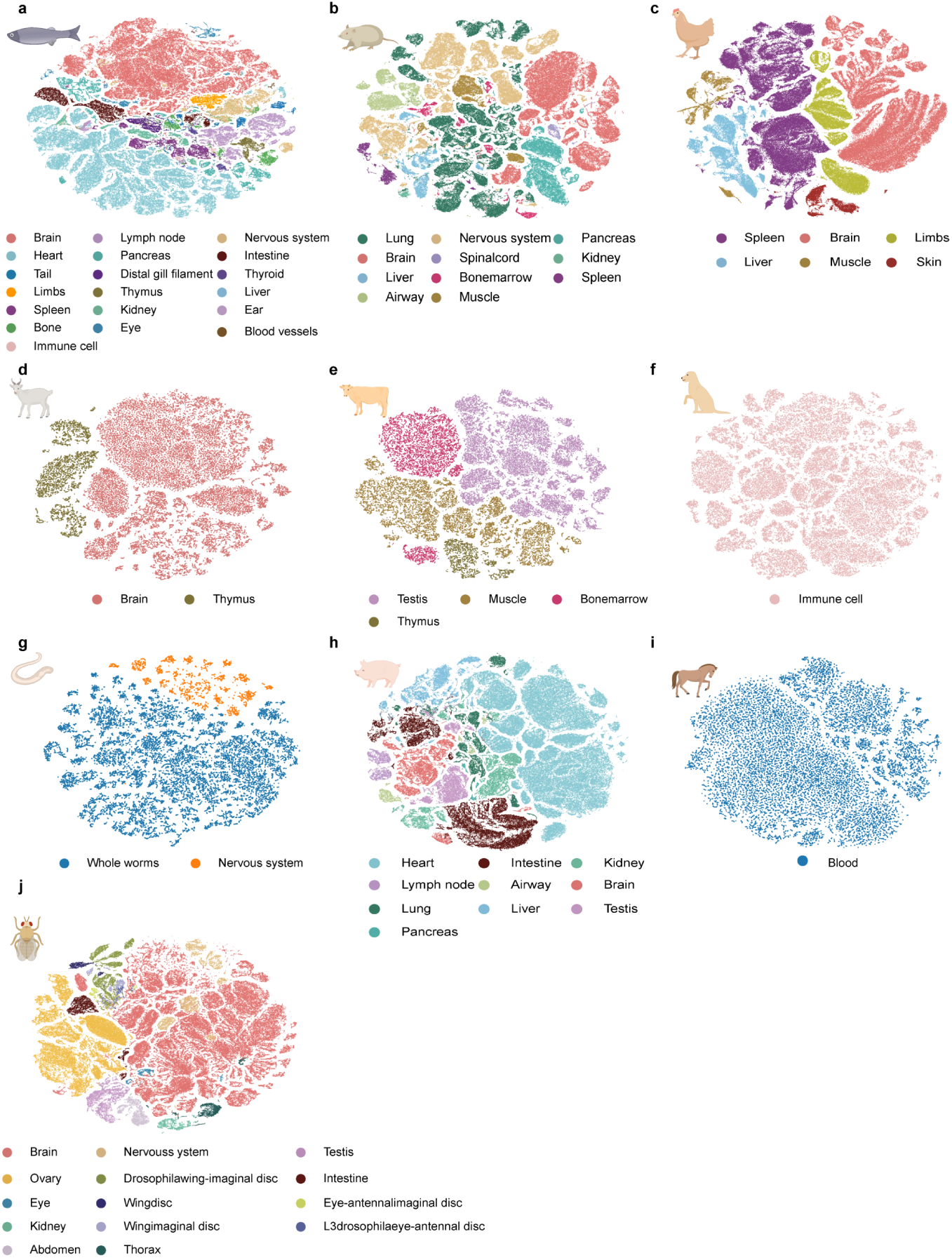
Single-cell Atlas Constrution of 11 species. tSNE plot showing single-cell atlas of Zebrafish, Fruit fly, Rat, Pig, Chicken, Bovine, Roundworm, Dog, Sheep, and Horse, dots with colors represent different tissues.

**Fig. S3.**
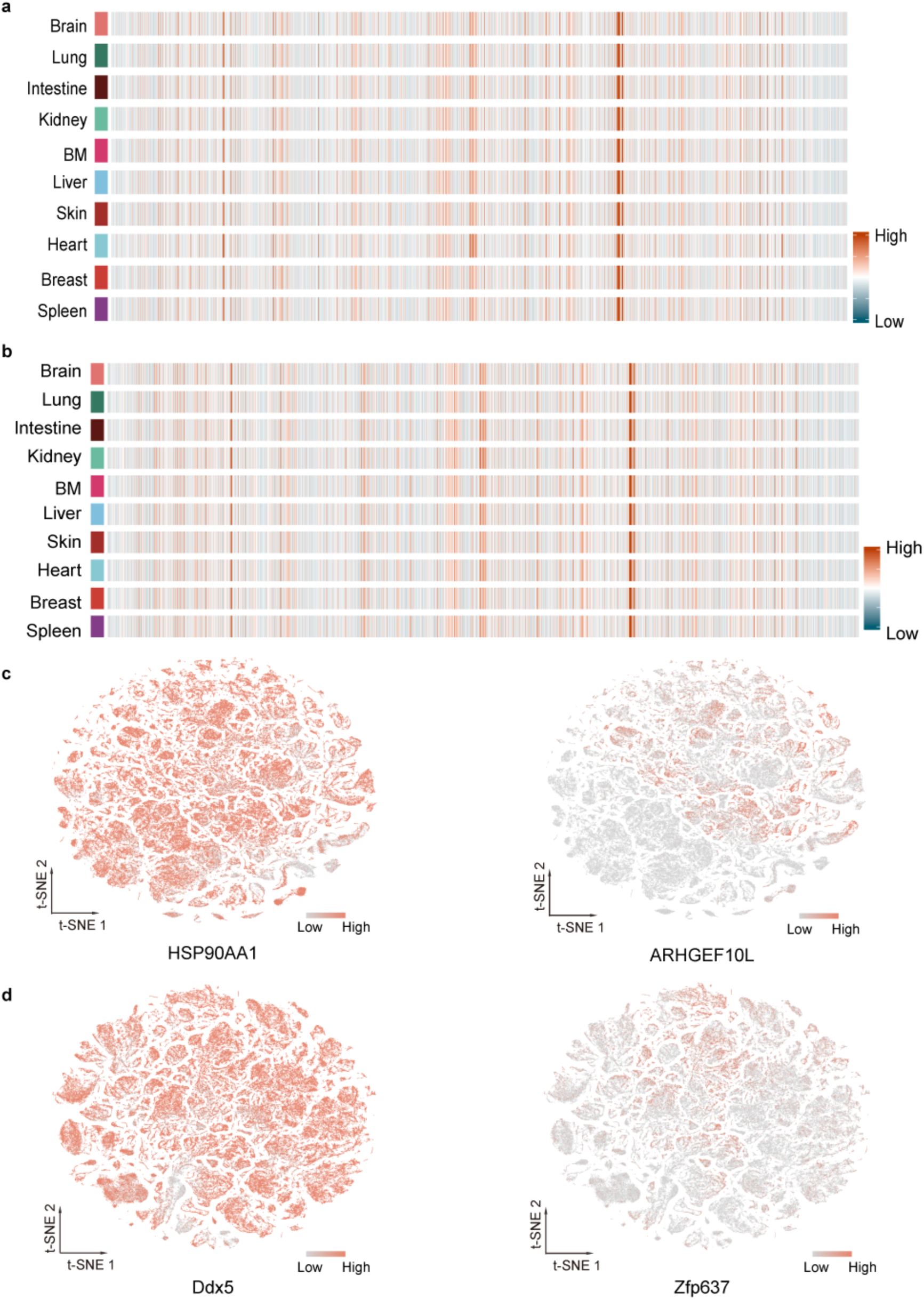
SEG analysis of human and mouse. Heatmap showing the distribution of HKGs in the top 10 organs for human (**a**) and mouse (**b**). **c.** tSNE visualization showing the expression pattern of HSP90AA1 of unique hSEGs and ARHGEF10L of unique hHKGs, respectively, in indicated the top30 organs of human. **d,** tSNE visualization showing the expression pattern of Ddx5 of unique mSEGs and Zfp637 of unique mHKGs, respectively, in indicated the top30 organs of mouse.

**Fig. S4.**
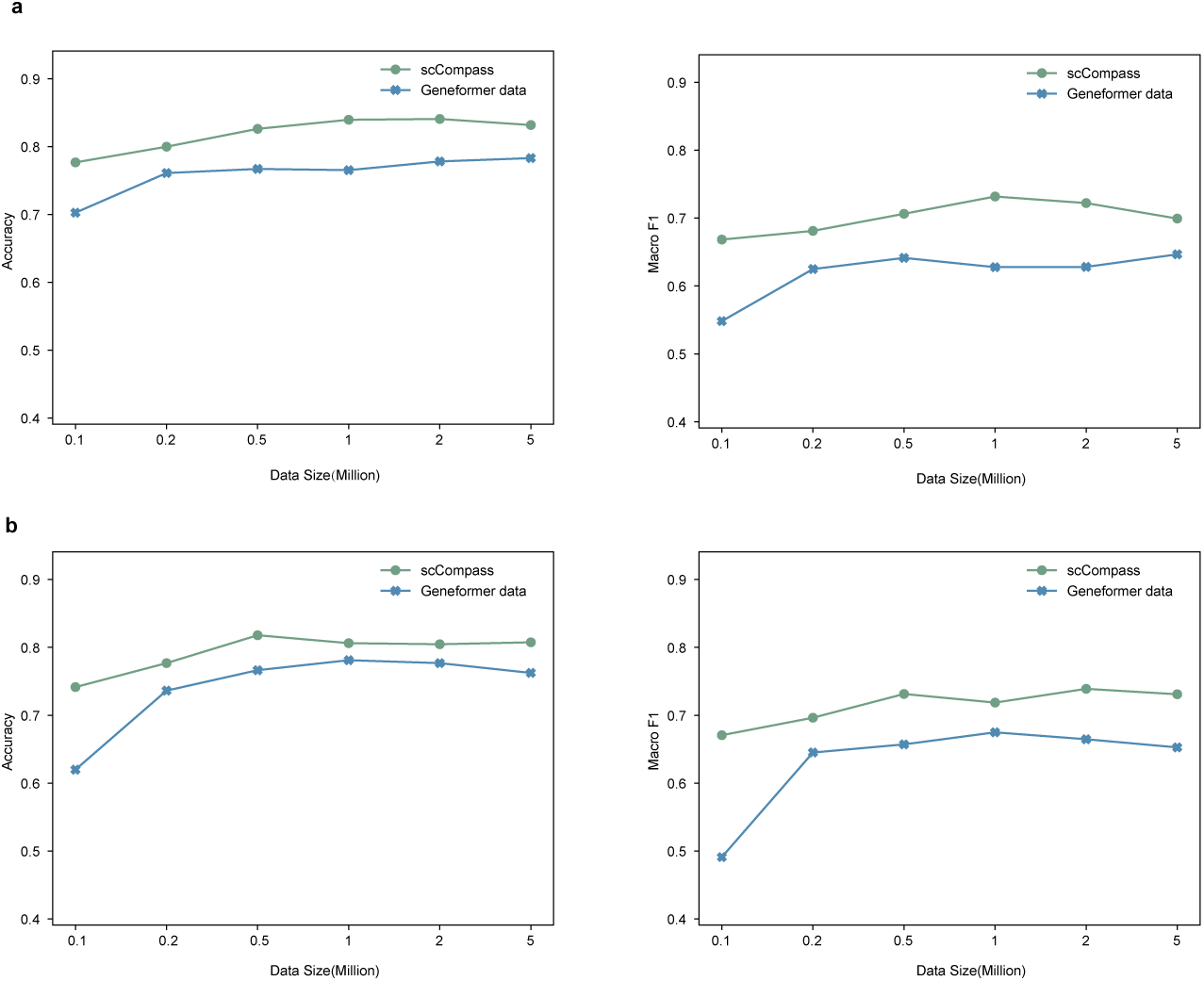
Evaluation of scCompass AI-Ready adaptability. Accuracy and Macro F1 of GeneFormer models trained on various scales of single-cell samples, evaluated on hMS and hLiver for cell type annotation. Line plots, with green points representing scCompass pretraining and blue points for Geneformer.

## Data and code availability

All primary data presented in this study will be deposited in a public database, and all codes including data preprocessing, model pre-training, multiple downstream tasks fine-tuning and the corresponding examples were uploaded to GitHub repository.

## Author Contributions

Yuanchun Zhou, Xin Li, Guihai Feng, Xuezhi Wang, Zhen Meng, and Shihua Zhang supervised the project. Yuanchun Zhou, Xin Li, Guihai Feng, Xuezhi Wang, Zhen Meng, Pengfei Wang, and Jiajia Wang conceived and designed the study. Pengfei Wang, Wenhao Liu, Jiajia Wang, Yana Liu, Pengjiang Li, Ping Xu, Wentao Cui, Ran Zhang, Qingqing, Jingxi Dong, Chunyang Zhang, Chengrui Wang and Guole Liu involved in collecting and preprocessing scRNA-seq data, designing the algorithm and performed the experiments. Pengfei Wang, Wenhao Liu, Jiajia Wang, and Yan Chen designed the interactive system. Pengfei Wang, Wenhao Liu, Jiajia Wang, Yana Liu and Pengjiang Li wrote the manuscript. Zhilong Hu, Chen Fang, Hanyu Xie, Yiyang Zhang, Meng Xiao, and Shubai Chen provided assistance in data collection and analysis. The X-COMPASS Project Consortium members collaborated in the paper discussions. All authors read and approved the final version of the manuscript.

## Acknowledgments

This research was funded by the National Natural Science Foundation of China (92470204), and the Informatization Plan of Chinese Academy of Sciences (CAS-WX2021SF-0101)

## APPENDIX

### The X-Compass Consortium Members

#### Institute of Zoology, Chinese Academy of Sciences

Xin Li, Hongmei Wang, Baoyang Hu, Wei Li, Fei Gao, Jingtao Guo, Leqian Yu, Qi Gu, Weiwei Zhai, Zhengting Zou, Guihai Feng, Wenhao Liu, Yao Tian, Chen Fang, Jingxi Dong, Yana Liu, Jingqi Yu, Wenhui Wu, Stella Lin, Cong Li, Yu Zou, Yongshun Ren, Fan Li, Yixiao Zhao, Yike Xin, Longfei Han, Shuyang Jiang, Kai Ma, Qicheng Chen, Haoyuan Wang, Huanhuan Wu, Chaofan He, Yilong Hu, Shuyu Guo, Yiyun Li

#### Computer Network Information Center, Chinese Academy of Sciences

Yuanchun Zhou, Yangang Wang, Xuezhi Wang, Pengfei Wang, Fei Li, Zhen Meng, Zheng Li, Zaitian Wang, Ping Xu, Wentao Cui, Zhilong Hu, Yang Wang, Huimin He, Shan Zong, Jiajia Wang, Yan Chen, Chunyang Zhang, Chengrui Wang, Qingqing Long, Ran Zhang, Meng Xiao, Qinmeng Yang, Zijian Wang, Yining Wang

#### Institute of Computing Technology, Chinese Academy of Sciences

Yiqiang Chen, Yi Zhao, Xiaodong Yang, Dechao Bu, Xin Qin, Jiaxin Qin, Zhaohui Yang, Chenhao Li, Zhufeng Xu, Zeyuan Zhang, Xiaoning Qi, Shubai Chen, Wuliang Huang, Yaning Li

#### Institute of Automation, Chinese Academy of Sciences

Ge Yang, Jing Liu, Guole Liu, Jie Jiang, Xingjian He, Liqun Zhong, Yaoru Luo, Jiaheng Zhou, Zichen Wang, Qinxuan Luo, Ziwen Liu, Ao Li, Teng Wang, Yiming Huang, Handong Li

#### Academy of Mathematics and Systems Science, Chinese Academy of Sciences

Yong Wang, Shihua Zhang, Jiahao Zhang, Yiyang Zhang, Shirui Li, Zhongming Liang, Zhenpeng Man, Kangning Dong, Qunlun Shen

